# Individual Differences in Sustained Attention Tune Semantic Abstraction in Neural Encoding of Narratives

**DOI:** 10.64898/2026.10.21.760815

**Authors:** Shan Gao, Yuan Chang Leong, Monica D. Rosenberg

## Abstract

Neural encoding modeling for narrative comprehension—using LLM embeddings to predict the temporal dynamics of neural activation—is widely used to evaluate how well the brain represents certain semantic information. This approach, however, typically collapses semantic representation into a scalar fidelity score aggregated across participants, overlooking potentially rich individual differences and qualitative differences in naturalistic neural representations. In the current study, we addressed this gap by investigating how neural semantic encoding varies across individuals with sustained attention, and whether this difference is qualitative in addition to quantitative. Using fMRI data from 49 participants listening to an audiobook, *Le Petit Prince*, in English, we trained encoding models with GPT-2 embeddings at varying layers and forecasting lengths. Encoding models trained on high-level embeddings (from the 8th layer of GPT-2) showed significant brain-LLM alignment in 73 of 122 cortical and subcortical ROIs, concentrated in the temporal cortex and the default mode network. Functional connectivity strength in a validated connectome-based predictive model of sustained attention correlated with encoding performance in 18 ROIs, most strongly in the insula and in control and attention networks, with no ROI showing significant opposite relationship. Decomposing semantic information by levels of abstraction showed that the neural signature of sustained attention was selectively associated with a greater proportion of high-level gist information encoded in the frontoparietal and attention networks, while the opposite relationship emerged for proportion of low-level information. Sustained attention scores further predicted stronger predictive coding of upcoming narrative content across multiple networks across the whole brain (control, default mode, salience/ventral attention, limbic, and somatomotor networks). These findings suggest that sustained attention does not uniformly amplify neural semantic encoding, but instead shifts neural representations toward more compressed, gist-level formats that support anticipation of upcoming content. Individual differences in brain-LLM alignment can be a signature of representational strategy rather than noise, with potential utility for cognitive phenotyping.

---

Narrative comprehension has become a powerful testbed for studying how the brain constructs meaning under ecologically valid conditions. Unlike understanding isolated words or sentences, comprehending narratives requires continuous integration of semantic information across multiple timescales, from individual words to unfolding event structures (Hasson et al., 2015). Together with the use of naturalistic stimuli, the use of large language model (LLM) embeddings as a computational proxy for time-resolved semantics has enabled researchers to quantify whether, and how strongly, different brain regions dynamically track semantic contents during narrative comprehension (Caucheteux et al., 2023; Goldstein et al., 2025; Tikochinski et al., 2025).

Within this paradigm, individual differences in neural encoding performance—measured as the correlation between predicted and ground truth neural dynamics—are typically treated as a noise to be ignored rather than a variable of theoretical interest; standard practice relies on group-level encoding models or reporting (e.g., Caucheteux et al., 2023; Tikochinski et al., 2025). A small number of recent studies in the broader field of neural encoding have begun to challenge this convention by linking trait-level cognitive phenotypes to individual differences in encoding performance. For example, work has reported associations between autism and atypical patterns of audiovisual stimulus encoding (Vanneau et al., 2025), between linguistic ability, attention, and encoding strength (Gu et al., 2025), and between various cognitive scores and inter-subject neural synchronization as well as language and visual feature encoding (Williams & Wehbe, 2021). Yet the relationship between attention and semantic encoding is not straightforward: in Gu et al. (2025), attentional orienting and executive control—as measured by the Attention Network Test—correlated with encoding performance in second-language speakers but not in first-language speakers. Rather than a simple "better attention, stronger encoding" relationship, this pattern implies that attention’s role may be context- or population-dependent. This raises an open mechanistic question: what, precisely, is attention doing to semantic encoding when it does matter?

Part of the difficulty in answering this question may lie in how encoding quality itself is operationalized. Existing approaches largely reduce encoding quality to a single scalar value; typically, the correlation between ground truth neural dynamics and the neural dynamics predicted from an embedding layer (Antonello et al., 2023; Caucheteux et al., 2023). This treats semantic encoding as a one-dimensional construct: better or worse, but not different in kind. By design, this approach cannot distinguish between two conceptually distinct possibilities: (1) that the brain encodes more or less semantic contents overall, versus (2) that it encodes semantic content at a different level of abstraction, ranging from local, lexical-level details to more global, gist-level latents. Looking at representational format complements and extends existing working linking attention to semantic encoding via scaling a fidelity metric up/down (Corriveau et al., 2026).

We propose a conceptual reframing that addresses this gap. Rather than uniformly increasing semantic encoding strength, we hypothesize that sustained attention improves encoding efficiency by prioritizing semantic compression during narrative comprehension: shifting representations from lower-level semantic details toward higher-level, compressed "gist" representations. This reframes individual differences in sustained attention from a "volume knob" model, in which attention simply amplifies a fixed representational format, to a "representational format" model, in which attention determines what kind of representation the brain constructs in the first place.

To test this account, we utilized a key property of deep neural networks: representations from different layers correspond to different levels of semantic abstraction, from local lexical details in early layers to more global narrative information in middle-later layers (Jawahar et al., 2019; Johnson et al., 2016; Kim et al., 2025). This layer-wise decomposition allows brain-LLM alignment to be computed separately for different levels of semantic information, rather than collapsed into a single fidelity score. This is the key methodological move that operationalizes our conceptual claims: it reframes the question from *how strongly* brain and LLM representations align to *what information* is encoded in the brain. Here, LLM embeddings not only serve as an encoding predictor, but also as a theoretical tool for probing the representational hierarchy of the brain during narrative comprehension.

Representational level is not merely descriptive; it carries functional consequences for prediction. Lower-level semantic details alone are insufficient for anticipating future content, whereas gist-level abstraction may be necessary to generate useful predictions about what comes next (Baldassano et al., 2017, 2018). For example, abstracting away from the colors and movements in a series of movie scenes to the gist of “taking a flight” enables us to predict “walking to the gate” following “going through security”, whereas passively mirroring the low-level visual details moment-by-moment does not help us anticipate what the main character will do next. If sustained attention indeed promotes this shift toward gist-level compression, we would further expect individuals with higher sustained attention to show enhanced predictive coding of future narrative contents.

If gist-level representations indeed support anticipation of upcoming narrative content, this predictive coding should, in principle, be directly measurable in neural dynamics rather than inferred solely from representational level. A complementary line of work has approached this by testing whether encoding models can be extended forward in time: rather than aligning neural responses to current semantic contents, this line of work asks whether neural dynamics also carry information about future semantic contents (Caucheteux et al., 2023; Goldstein et al., 2022). Because upcoming and current content are themselves auto-correlated, isolating a genuine predictive signal requires removing the semantic information already encoded in the current semantic embeddings; in this study, we regressed out current embeddings from future embeddings. This yields a measure of predictive coding conceptually distinct from standard, same-timepoint encoding performance. If sustained attention promotes a shift toward more abstract, compressed representations, this shift should manifest not only as a change in representational level but also as a measurable gain in the brain’s capacity to predictively encode future narrative contents.

We tested the above hypotheses and framework using the *Le Petit Prince* fMRI dataset (Li et al., 2022). The amount of semantic information encoded in neural dynamics was operationalized as encoding model performance, using GPT-2 embeddings as predictors. Individual-specific sustained attention was estimated using the predictions of a connectome-based predictive model of sustained attention (saCPM) trained on an independent dataset and population (Rosenberg et al., 2016). We first show that LLM embeddings can predict neural dynamics, replicating the brain-LLM semantic alignment shown in prior work (Caucheteux & King, 2022; Goldstein et al., 2022). We then show that individual-specific sustained attention estimates correlate with semantic encoding strength in general. We go on to show that these attention estimates correlate specifically with the balance between local and gist-level semantic representation, consistent with a compression account, and we connect this representational shift to enhanced predictive coding of upcoming content. Together, these findings reframe individual differences in narrative encoding as a source of theoretical insights into the computational properties of language processing rather than noises to be averaged away. Individual differences in sustained attention predict qualitative differences in representational strategy in addition to quantitative differences in representational capacity.

## Methods

### fMRI dataset

We utilized a publicly available fMRI dataset by Li et al. (2022). Forty-nine English speakers listened to the audiobook *Le Petit Prince* in English while undergoing fMRI scans. The narrative was divided into nine consecutive sections, each was a separate run and lasted approximately 10 minutes. Functional images were acquired with a repetition time (TR) of 2 s.

We used the preprocessed fMRI data provided by the original authors. Steps included correction for slice-timing differences (3dTshift), despike (3dDespike), volume registration (3dvolreg), anatomical alignment, nonlinear alignment to the MNI template brain, denoising with multi-echo independent components analysis (ME-ICA) for motion, physiology and scanner artifacts, resampled at 2 mm cubic voxels (3dresample) (Li et al., 2022). Following prior work on attentional engagement during narrative comprehension (Song et al., 2021), we parcellated the data into 114 cortical regions of interest (ROIs) using the Yeo atlas (Yeo et al., 2011) and 8 subcortical ROIs using the Brainnetome atlas (Fan et al., 2016), resulting in a total of 122 ROIs for each participant. For each participant and ROI, we extracted the mean BOLD time series.

### LLM semantic representations

To acquire moment-by-moment semantic representations of the narrative in LLMs, we extracted contextualized token embeddings from the 8th layer of GPT-2 model (768 dimensions), which has been shown to capture high-level semantic information and align with human neural representations of narratives (Caucheteux et al., 2023). The tokenized transcript was processed using a sliding context window of up to 1024 tokens preceding each token (inclusive of current token), the largest context window size of GPT-2.

Token-level embeddings were resampled to TR-level by aggregating all tokens whose timestamps fell within each TR (number of tokens per TR: *M* = 7.31, *SD* = 1.66). For each TR, we computed a single semantic representation by mean-pooling the embeddings of all tokens in that interval, yielding a TR-by-feature matrix.

### Measuring brain-LLM alignment with encoding models

We operationalized brain–LLM alignment using encoding models. For each participant and each ROI, we trained an encoding model to predict the ROI’s BOLD time course from LLM-derived semantic representations.

The encoding pipeline consisted of principal component analysis (PCA) for dimensionality reduction followed by ridge regression. PCA was first applied to reduce the dimensionality of LLM-derived semantic representations to 32 principal components following prior work (Caucheteux et al., 2023). Then, a ridge regression was fitted on the 32 principal components to predict neural dynamics. The regularization parameter (α) for each encoding model was optimized via nested five-fold cross-validation over a logarithmically spaced range from 10⁻¹ to 10⁸.

Encoding models were trained on sections 1∼6 and evaluated on held-out sections 7∼9. To account for the hemodynamic lag, we shifted fMRI time series by 2 TRs (4 s) relative to the semantic predictors. For each ROI, encoding performance was evaluated with Pearson’s correlation between the predicted and observed neural dynamics in the test set.

Statistical significance of encoding performance was assessed using a permutation-based null distribution. For each participant and ROI, the observed BOLD time course in the test set was phase-randomized 1,000 times, and a null encoding performance was computed for each permutation. For this analysis, both true and null encoding performance were aggregated across participants by taking the median. The actual encoding performance was significant if it exceeded the 95th percentile of the null distribution and survived *FDR* correction (*q* < .05). All analyses in this study (including the ones described below) were run with a random seed 1234.

### Connectome-base predictive model (CPM) of individual sustained attention

To estimate individual differences in sustained attention, we applied a pretrained connectome-based predictive model, the sustained attention CPM (saCPM; Rosenberg et al., 2016) to fMRI data from sections 1∼6. The saCPM was trained using data from Rosenberg et al. (2016) and nodes in Song et al. (2021). For each participant, a functional connectivity matrix was computed using ROI-level BOLD time courses across the six sections (1810 TRs in total).

The saCPM was trained on a separate group of participants performing the gradual-onset continuous performance task (gradCPT; Rosenberg et al., 2016). The trained CPM was then applied to the present dataset to generate an individual sustained attention score for each participant: functional connectivity values in each network mask were averaged and the difference in mean connectivity in the positive mask (which predicts better sustained attentional performance) and negative mask (which predicts worse performance) was taken.

### Relating brain-LLM alignment to the sustained attention CPM

To examine the relationship between individual differences in brain–LLM alignment during narrative comprehension and the neural signature of sustained attention, we computed, for each ROI, a rank-order correlation (Spearman’s *r*) across participants between encoding performance and sustained attention scores (**Figure 1**).

**Figure 1.**
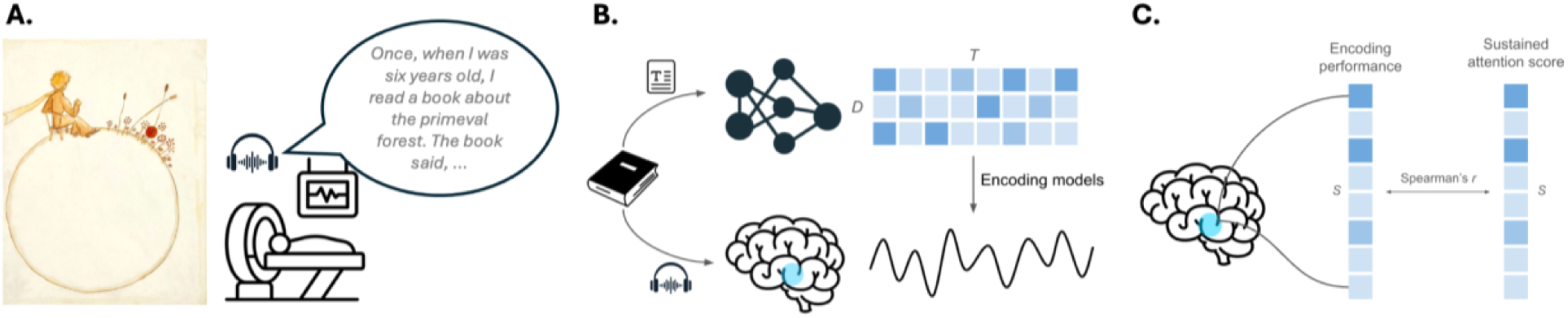
Measuring brain-LLM alignment and its relationship to sustained attention. **A.** We used a publicly available fMRI dataset by Li et al. (2022), where 49 English speakers listened to the audiobook *Le Petit Prince* while undergoing fMRI scans. **B.** We operationalized brain–LLM alignment using encoding performance. For each participant and each ROI, we trained an encoding model to predict the ROI’s BOLD time course from LLM-derived semantic representations. **C.** For each ROI, we related individual differences in brain–LLM alignment to sustained attention by computing their rank-order correlation across participants.

Statistical significance was assessed using a two-tailed permutation test where participant labels were randomly shuffled 1,000 times. For each permutation, the rank-order correlation was recomputed, generating a null distribution against which the observed correlation was compared.

### Decomposing high-level and low-level semantic information

Low-level and high-level semantic representations were operationalized using layer 1 and layer 8 embeddings, respectively, from the GPT-2 model using the same protocol as before (maximum 1024 token context windows; converting per-token embeddings to per-TR embeddings). To isolate the unique contribution of high-level semantic information, we orthogonalized the two embedding sets. Layer 1 embeddings were first reduced to 32 principal components. We then regressed layer 8 embeddings onto these layer 1 components using ordinary least squares (OLS) regression and retained only the residuals; these residuals were finally reduced to 32 principal components as well to be used as the predictors of neural encoding models. This removes shared variance between the two sets of embeddings so that layer 8 components capture isolated high-level semantics whereas layer 1 components capture low-level semantics.

Encoding models were fit with both embedding sets, respectively, for all participant and ROI combinations. Encoding model fitting followed the same procedure as described in previous sections (ridge regression; regularization optimized via nested five-fold cross-validation; trained on story sections 1–6 and evaluated on held-out sections 7–9). Encoding performance was quantified as the correlation between predicted and observed neural dynamics in the test set. Group-level statistical significance of encoding performance was assessed by comparing the median performance across participants against a null distribution generated from 1,000 phase-randomized permutations. We denote the encoding performance using layer 8 components as *encoding_hi_*, and the encoding performance using layer 1 components as *encoding_lo_*.

Because we are interested in the degree of semantic compression in the brain, the *proportion* of high-level information encoded relative to low-level information in addition to an absolute *encoding_hi_* value. For each ROI and each participant, we computed proportion of high-level information encoded as follows:

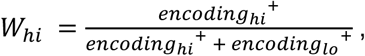

where each encoding performance was thresholded at 0 to preserve interpretability of proportion.

To test whether the proportion of high-level information encoded in each ROI scales with sustained attention, we ran analyses on both network and per-ROI level. At network level, we fit a mixed effect linear regression model for each network:

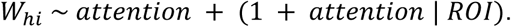

At per-ROI level, we computed the rank-order correlation between individual-specific sustained attention scores and *W_hi_*; two-tailed statistical significance was assessed by a null distribution generated by 1,000 random permutations of attention score ordering. As validation analyses, we also tested the relationship between individual-specific raw *encoding_hi_* scores and sustained attention scores using the same approach.

Additionally, to further test the specificity of any patterns found to high-level information encoding, we ran control analyses by isolating low-level semantics, i.e., regressing out layer 8 variances from layer 1 embeddings, and repeating the analyses described above.

### Measuring predictive coding in the brain

To measure how well the brain predictively encodes upcoming story content, we isolated moment-to-moment future information from GPT-2 layer 8 embeddings and measured how well it predicted neural dynamics.

Different from previous analyses, per-token embeddings were extracted using a context window of maximum 128 tokens instead of 1024 tokens to surface some temporal dynamics in semantic information given the high autocorrelation of *Le Petit Prince*. For each TR, a raw future embedding was constructed by averaging the token embeddings spanning the 7 tokens preceding the forecast horizon (25 tokens ahead), and a current embedding was constructed using the same protocol as before (i.e., averaging the token embeddings within the current TR). To isolate the unique contribution of future information, we regressed out current information from raw future embeddings: first, reducing the current embeddings to 32 principal components via PCA; then orthogonalizing the raw future embeddings against them by regressing the current components out via OLS, retaining only variance unique to future information. The residualized future embeddings were then reduced to 32 principal components.

Using ridge regression with the regularization penalty tuned by nested five-fold cross-validation, we fit encoding models for each participant and ROI combination with residualized future embeddings. Models were fit on training sections (1–6) and evaluated on held-out sections (7–9). Predictive coding performance was quantified as the correlation between predicted and observed neural dynamics. Group-level statistical significance was assessed by comparing the median performance across participants against a null distribution generated from 1,000 phase-randomized permutations of the test timecourses.

We tested whether predictive coding performance scales with neural signatures of sustained attention at both network and per-ROI level. At the network level, we fit a mixed effect linear regression model for each network:

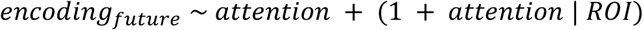

The same regression model was also fit on ROIs across the whole brain to detect any global effect. At per-ROI level, we computed rank-order correlation between individual-specific saCPM scores and predictive coding performance, and assessed two-tailed statistical significance by a null distribution generated by 1,000 random permutations of attention score ordering.

## Results

### Semantic representations in brain and LLM align during narrative comprehension

We first assessed whether brain and LLM semantic representations align during narrative comprehension. Encoding models were trained to predict ROI-level neural dynamics from GPT-2 semantic representations and evaluated on held-out sections.

As shown in **Figure 2**, across all ROIs, 73 out of 122 showed significant encoding performance. The highest brain-LLM alignment was observed in the temporal lobe, consistent with its established role in speech and narrative comprehension. Significant brain-LLM alignment was also observed across the default mode network (DMN). Together, these results support shared semantic representations in the brain and LLM.

**Figure 2.**
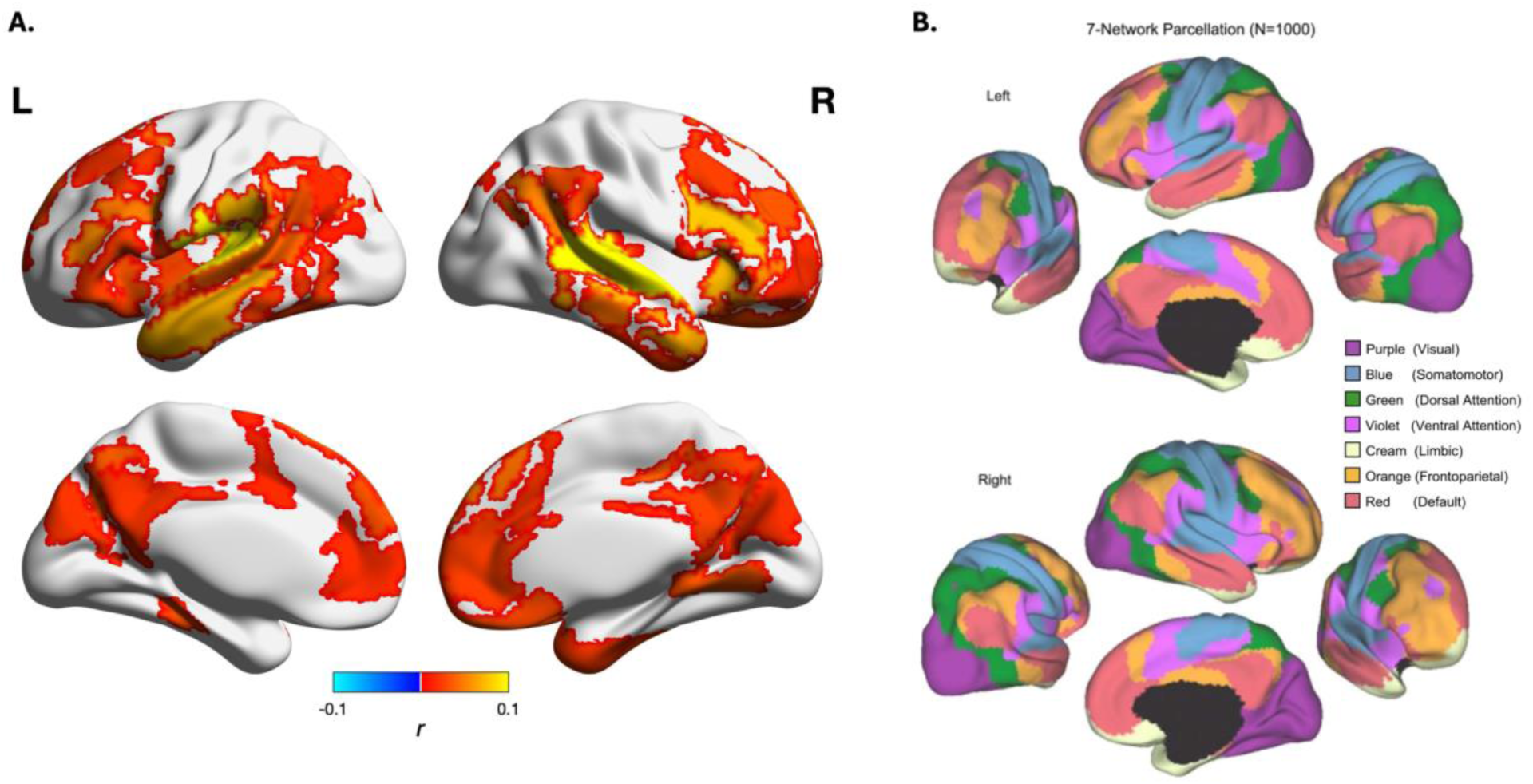
**A**. Performance of individual-specific encoding models, aggregated across participants by taking the median (FDR *q* < .05). The predictors of the encoding models were first reduced to 32-dimension with PCA. Then a ridge regression was fit for each participant and ROI, with optimized alpha (logspace 10^−1^∼10^8^) selected through a nested 5-fold cross-validation. Statistical significance was assessed by phase-randomizing ground-truth brain time courses in the test set 1,000 times per participant and ROI combination; for each ROI, a null mean performance was computed each iteration by taking the median of null performance across participants. **B**. An illustration of 7-network parcellation from Yeo et al. (2011).

### Individuals with neural signatures of stronger sustained attention show greater brain-LLM alignment

We then examined whether a neural signature of sustained attention was related to the degree of brain–LLM alignment during narrative comprehension. Individual-specific sustained attention scores were estimated using a pretrained connectome-based predictive model (CPM) applied to functional connectivity during the narrative.

At the network level, significant correlation between encoding performance and sustained attention was strongest in the control and attention networks, including Control Network A (*r* = 0.25), Control Network B (*r* = 0.24), Dorsal Attention Network B (*r* = 0.28), Salience/Ventral Attention Network A (*r* = 0.38), Salience/Ventral Attention Network B (*r* = 0.27), and Somatomotor Network B (*r* = 0.23).

Across all ROIs, 18 out of 122 (14.75%) showed a significant correlation between encoding performance and sustained attention estimates (**Figure 3A**). When restricting the analysis to ROIs that exhibited significant encoding performance, 13 out of 73 ROIs (17.81%) showed a significant correlation. In these ROIs, individuals with higher sustained attention scores tended to exhibit stronger brain-LLM alignment; none of the significant ROIs showed the opposite pattern. The strongest correlation between encoding performance and sustained attention was observed in the right insular cortex (*r* = 0.61). This is consistent with the insular cortex’s role as a hub for integrating emotional responses and story content (Nguyen et al., 2016), an essential part of engaging with a story.

**Figure 3.**
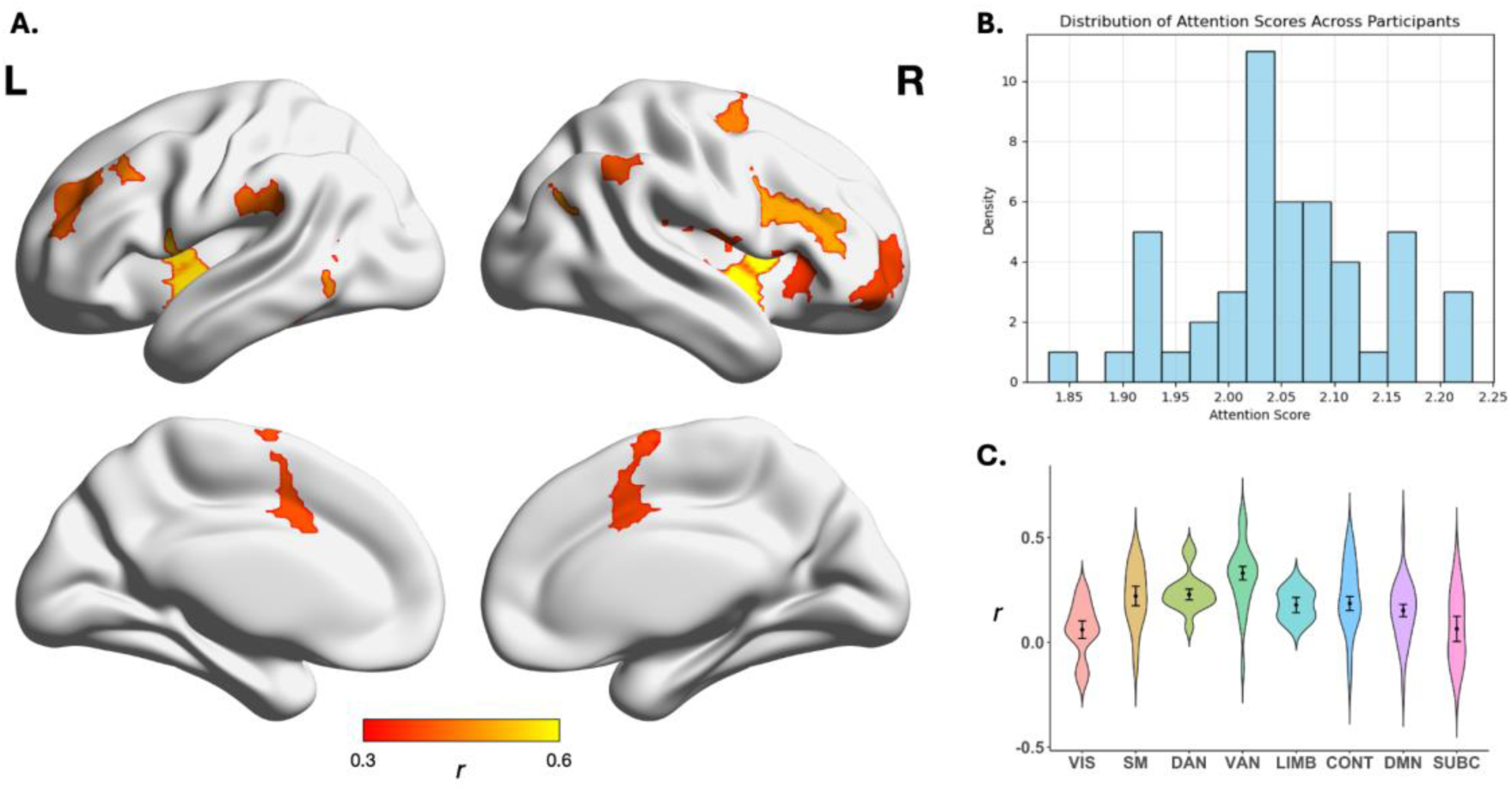
**A**. 18 ROIs with better encoding performance for participants with a higher sustained attention score. Color indicates Spearman’s correlation between sustained attention score and encoding performance across participants (FDR *q*-masked). Two-tailed statistical significance was tested by shuffling the order of participants 1,000 times. The highest correlation was observed in the insular cortex. **B**. Distribution of individual-specific sustained attention scores. The X-axis denotes the rank-order correlation between individual-specific sustained attention scores and neural encoding performance. **C**. The median and distribution of correlations between sustained attention score and encoding performance within each network.

### Semantic abstraction scales with a neural signature of sustained attention

How is higher neural encoding efficiency achieved with better sustained attention? We hypothesize that the brain actively infers the story’s high-level gist while processing narratives in real time, and that better sustained attention facilitates latent inference and therefore semantic compression.

We operationalized low-level semantics as the layer 1 embeddings of GPT-2 model, and high-level latent as the layer 8 embeddings of GPT-2 model regressed out of the variances in layer 1 embeddings. For each participant, we fit separate encoding models predicting per-ROI neural dynamics from low-level semantics and high-level latent respectively, and evaluated the models on held-out sections. Out of 122 ROIs, 38 ROIs showed significant encoding performance of high-level semantics (FDR *q* < .05; **Figure 4A**).

**Figure 4.**
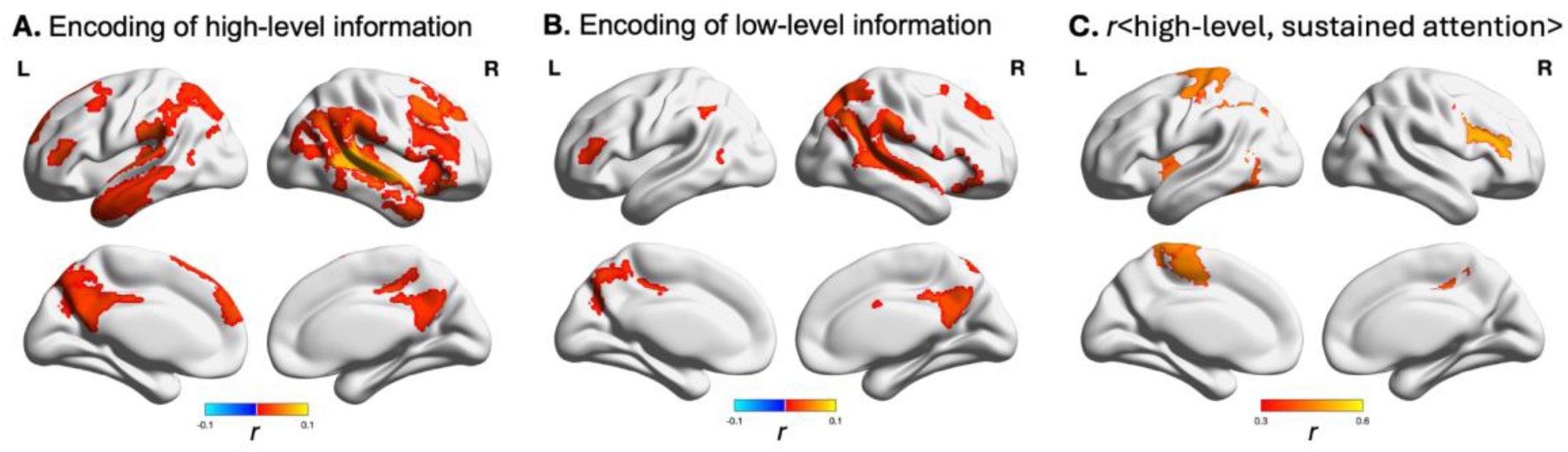
**A**. 38 out of 122 ROIs showing significant encoding of decomposed high-level semantic information (two-tailed; FDR *q* < .05). Decomposed high-level semantic information was constructed using the 8th layer embeddings of GPT-2 model regressed out of the variances of 1st layer embeddings. Results were aggregated across participants by taking the median. **B**. Decomposed low-level semantic information was significantly encoded in 18 out of 122 ROIs (two-tailed; FDR *q* < .05). Decomposed low-level semantic information was constructed using the 1st layer embeddings of GPT-2 model regressed out of the variances of 8th layer embeddings. Results were aggregated across participants by taking the median. On per-ROI level, no ROI showed significant correlation between *W_lo_* and sustained attention, or between *encoding_lo_* (encoding of low-level semantic information: 1st layer embeddings of GPT-2 model regressed out of the variances of 8th layer embeddings) and sustained attention (two-tailed; FDR *q* < .05). **C**. ROIs with significant correlation between *encoding_hi_* (encoding of high-level semantic information: 8th layer embeddings of GPT-2 model regressed out of the variances of 1st layer embeddings) and sustained attention (two-tailed; FDR *q* < .05). On per-ROI level, no ROI showed significant correlation between *W_hi_* and sustained attention (two-tailed; FDR *q* < .05).

We then tested whether the proportion of high-level information encoded in each ROI scales with sustained attention. The proportion of high-level information encoded was computed as high-level encoding performance divided by the sum of high- and low-level encoding performance, both thresholded at 0 to preserve interpretability of proportion. At network level, we fit a mixed effect linear regression model for each network:

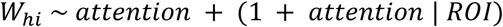

Across the 8 networks, the control network and the salience and ventral attention network showed significant positive relationship between sustained attention and proportion of high-level information (control network: *b* = 0.027, *SE* = 0.011, *z* = 2.376, *p* = .018, 95% CI [0.005, 0.049]; salience and ventral attention network: *b* = 0.030, *SE* = 0.014, *z* = 2.089, *p* = .037, 95% CI [0.002, 0.057]; **Table 1**). The salience and ventral attention network is involved in allocating cognitive resources to meaningful information and suppressing irrelevant information (Corbetta et al., 2008; Corbetta & Shulman, 2002); higher sustained attention may promote the focus on high-level information by making this gate function more effectively.

**Table 1.**
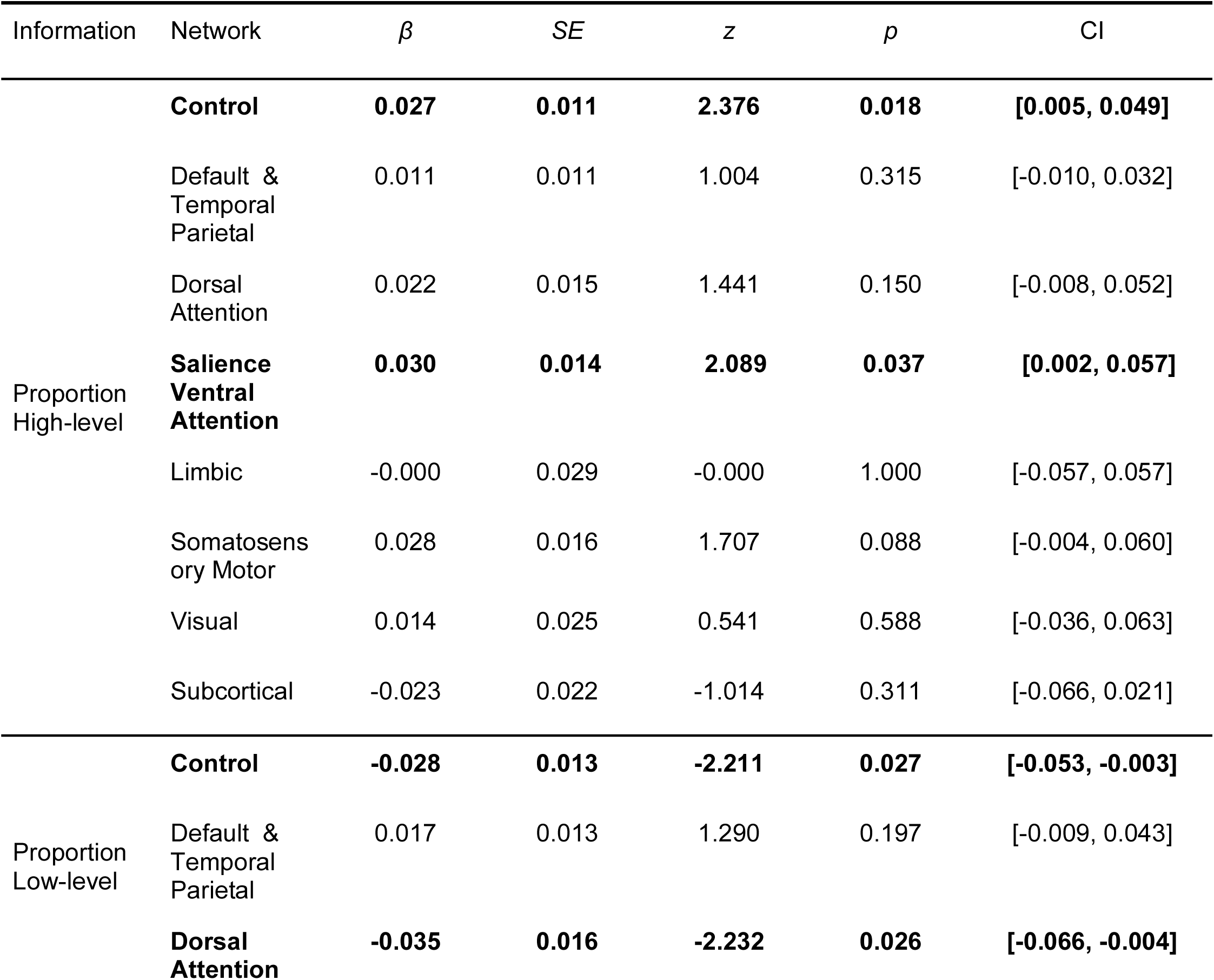
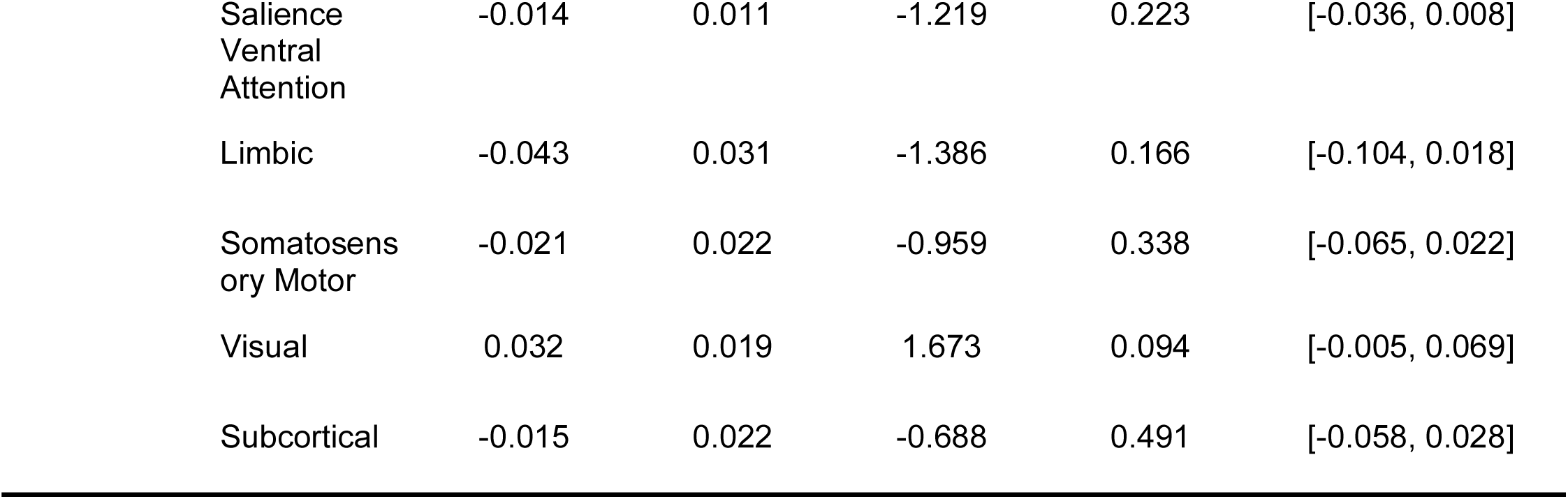
Network level mixed effect linear regression models predicting *proportion* of high-level and low-level semantic information (respectively) from sustained attention score. Proportion high-level information: Across the 8 networks, the control network and the salience and ventral attention network showed significant positive relationship between sustained attention and proportion of high-level semantic information. Proportion low-level information: Most networks showed a negative relationship between sustained attention and proportion of low-level information encoded, among which the control network and the dorsal attention network were significant.

| Information | Network | $\beta$ | $SE$ | $z$ | $p$ | CI |
| --- | --- | --- | --- | --- | --- | --- |
| Proportion High-level | **Control** | **0.027** | **0.011** | **2.376** | **0.018** | **[0.005, 0.049]** |
|  | Default & Temporal Parietal | 0.011 | 0.011 | 1.004 | 0.315 | [-0.010, 0.032] |
|  | Dorsal Attention | 0.022 | 0.015 | 1.441 | 0.150 | [-0.008, 0.052] |
|  | **Salience Ventral Attention** | **0.030** | **0.014** | **2.089** | **0.037** | **[0.002, 0.057]** |
|  | Limbic | -0.000 | 0.029 | -0.000 | 1.000 | [-0.057, 0.057] |
|  | Somatosensory Motor | 0.028 | 0.016 | 1.707 | 0.088 | [-0.004, 0.060] |
|  | Visual | 0.014 | 0.025 | 0.541 | 0.588 | [-0.036, 0.063] |
|  | Subcortical | -0.023 | 0.022 | -1.014 | 0.311 | [-0.066, 0.021] |
| Proportion Low-level | **Control** | **-0.028** | **0.013** | **-2.211** | **0.027** | **[-0.053, -0.003]** |
|  | Default & Temporal Parietal | 0.017 | 0.013 | 1.290 | 0.197 | [-0.009, 0.043] |
|  | **Dorsal Attention** | **-0.035** | **0.016** | **-2.232** | **0.026** | **[-0.066, -0.004]** |

**Table 1.** Network level mixed effect linear regression models predicting *proportion* of high-level and low-level semantic information (respectively) from sustained attention score.
|  |  |  |  |  |  |
| --- | --- | --- | --- | --- | --- |
| Saliency<br>Ventral<br>Attention | -0.014 | 0.011 | -1.219 | 0.223 | [-0.036, 0.008] |
| Limbic | -0.043 | 0.031 | -1.386 | 0.166 | [-0.104, 0.018] |
| Somatosensory<br>Motor | -0.021 | 0.022 | -0.959 | 0.338 | [-0.065, 0.022] |
| Visual | 0.032 | 0.019 | 1.673 | 0.094 | [-0.005, 0.069] |
| Subcortical | -0.015 | 0.022 | -0.688 | 0.491 | [-0.058, 0.028] |

On the per-ROI level, we computed rank-order correlation between individual-specific *W_hi_* values and sustained attention scores. No ROI stably showed as significant after FDR correction, indicating that the positive relationship between sustained attention and proportion of high-level information might be a small yet distributed effect in the above networks, which needs more statistical power to detect without pooling across ROIs.

To test if attention-enhanced semantic encoding is specific to high-level information instead of being universal across both high- and low-level information, we ran control analyses by isolating low-level semantics, i.e., regressing out layer 8 variances from layer 1 embeddings. Out of 122 ROIs, 18 ROIs showed significant encoding performance of low-level semantics (two-tailed; FDR *q* < .05; **Figure 4B**)—with smaller absolute values than high-level semantics (**Figure 4A**; **Figure 4B**).

We then computed proportion of low-level information encoded (*W_lo_*) in each ROI similarly to how we computed (*W_hi_*): low-level encoding performance divided by the sum of low- and high-level encoding performance, both thresholded at 0 to preserve interpretability of proportion. At the network level, we fit a mixed effect linear regression model for each network:

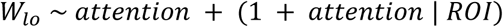

Contrary to high-level information, most networks showed a negative relationship between sustained attention and proportion of low-level information encoded, among which the control network and the dorsal attention network were significant (control network: *b* = −0.028, *SE* = 0.013, *z* = −2.211, *p* = .027, 95% CI [-0.053, −0.003]; dorsal attention network: *b* = −0.035, *SE* = 0.016, *z* = −2.232, *p* = .026, 95% CI [-0.066, −0.004]; **Table 1**). On the per-ROI level, we computed rank-order correlation between individual-specific *W_lo_* values and sustained attention scores. Similar to *W_hi_*, no ROI was significant after FDR correction.

To further examine whether the negative correlation between *W_lo_* and sustained attention scores were driven by an absolute reduction of low-level information with higher sustained attention scores or by greater increase of high-level information than low-level information, we ran the above analyses with raw semantic encoding performance. Network level regression analyses showed that both high- and low-level information’s raw encoding performance correlated positively with sustained attention scores in the majority of networks, with more networks significant and generally higher coefficients for high-level semantic encoding (**Table 2**). At per-ROI level, raw high-level semantic encoding performance showed significant positive correlation with sustained attention scores in several ROIs (**Figure 4C**), while low-level semantic encoding performance showed no significant ROI after FDR correction.

**Table 2.** Network level mixed effect linear regression models predicting the raw encoding performance of high-level and low-level semantic information (respectively) from sustained attention score. High-level information encoding: Across the 8 networks, 6 networks showed significant positive relationship between sustained attention and high-level semantic encoding. Low-level information encoding: Most networks showed a small to near-zero positive relationship between sustained attention and low-level semantic encoding, among which the default & temporal parietal network, 2 attention networks, and the visual network were significant. Most networks, except for the visual network, showed a tendency towards more increase in high-level semantic encoding performance with increased attention score than low-level semantic encoding performance.

| Information | Network | $\beta$ | $SE$ | $z$ | $p$ | CI |
| --- | --- | --- | --- | --- | --- | --- |
| High-level | **Control** | **0.010** | **0.002** | **5.889** | **0.000** | **[0.006, 0.013]** |
|  | **Default & Temporal Parietal** | **0.008** | **0.002** | **5.252** | **0.000** | **[0.005, 0.011]** |
|  | **Dorsal Attention** | **0.010** | **0.003** | **3.644** | **0.000** | **[0.005, 0.015]** |
|  | **Salience Ventral Attention** | **0.011** | **0.002** | **6.191** | **0.000** | **[0.007, 0.014]** |
|  | Limbic | 0.004 | 0.003 | 1.123 | 0.262 | [-0.003, 0.010] |
|  | **Somatosensory Motor** | **0.007** | **0.002** | **2.653** | **0.008** | **[0.002, 0.011]** |
|  | **Visual** | **0.008** | **0.002** | **3.265** | **0.001** | **[0.003, 0.013]** |
|  | Subcortical | 0.005 | 0.003 | 1.737 | 0.082 | [-0.001, 0.010] |
| Low-level | Control | 0.002 | 0.002 | 1.138 | 0.255 | [-0.001, 0.005] |
|  | **Default & Temporal Parietal** | **0.007** | **0.002** | **4.559** | **0.000** | **[0.004, 0.010]** |
|  | **Dorsal Attention** | **0.005** | **0.002** | **2.365** | **0.018** | **[0.001, 0.008]** |
|  | **Salience Ventral Attention** | **0.003** | **0.001** | **2.427** | **0.015** | **[0.001, 0.006]** |
|  | Limbic | -0.000 | 0.005 | -0.062 | 0.950 | [-0.011, 0.010] |
|  | Somatosensory Motor | 0.003 | 0.003 | 0.972 | 0.331 | [-0.003, 0.008] |
|  | **Visual** | **0.011** | **0.002** | **4.458** | **0.000** | **[0.006, 0.015]** |
|  | Subcortical | 0.003 | 0.002 | 1.355 | 0.175 | [-0.001, 0.008] |

The above evidence supports the hypothesis that sustained attention is associated with selective increases in high-level encoding—greater semantic abstraction—instead of simple global increase of encoding performance.

### Neural signature of sustained attention predicts stronger forecasting

Being aware of a story’s gist may enable the brain to predictively encode future contents of the story. Therefore, we tested whether the brain encodes future information, and whether predictive coding is associated with a neural signature of better sustained attention. Future information embeddings were constructed with mean future token embeddings up to 25 tokens ahead of time, regressed out of the variances in current token embeddings. Due to the high autocorrelation of *Le Petit Prince*, we shortened the context window size to 128 tokens when extracting embeddings from GPT-2, to surface some temporal dynamics in semantic information. Aggregating across participants (taking median), we found no ROI with significant predictive coding performance.

However, a non-significant predictive coding performance aggregated across participants may result from participants with lower sustained attention scores contributing low forecasting performance and does not rule out the potential relationship between semantic forecasting and sustained attention. Therefore, we then tested if participants with higher sustained attention scores had better predictive coding performance.

At network level, we fit a mixed effect linear regression model for each network:

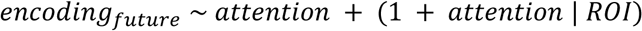

Across the 8 networks, 5 networks showed a significant positive relationship between sustained attention score and predictive coding (**Table 3**): the control network (*b* = 0.002, *SE* = 0.001, *z* = 1.988, *p* = .047, 95% CI [0.000, 0.005]), the default mode network (*b* = 0.005, *SE* = 0.001, *z* = 4.477, *p* = .000, 95% CI [0.003, 0.008]), the salience and ventral attention network (*b* = 0.004, *SE* = 0.001, *z* = 2.655, *p* = .008, 95% CI [0.001, 0.006]), the limbic network (*b* = 0.006, *SE* = 0.003, *z* = 2.150, *p* = .032, 95% CI [0.001, 0.012]), and the somatomotor network (*b* = 0.004, *SE* = 0.002, *z* = 2.075, *p* = .038, 95% CI [0.000, 0.007]). In addition to the networks relevant to attentional allocation, here we also found DMN significant. This may be due to DMN being essential for representing event schemas (Baldassano et al., 2018; Yeshurun et al., 2021), which is needed for predictive coding. The same mixed effect linear regression model run on ROIs across the whole brain showed the same pattern of positive relationship between sustained attention and predictive coding (*b* = 0.003, *SE* = 0.001, *z* = 6.356, *p* = .000, 95% CI [0.002, 0.004]).

**Table 3.** Network level mixed effect linear regression models predicting encoding of future content from sustained attention score. Most networks showed a positive relationship, among which 5 were significant: the control network, the default mode network, the salience/ventral attention network, the limbic network, and the somatomotor network.

| | $\beta$ | $SE$ | $z$ | $p$ | CI |
| --- | --- | --- | --- | --- | --- |
| **Cont** | **0.002** | **0.001** | **1.988** | **0.047** | **[0.000, 0.005]** |
| **Default & TempPar** | **0.005** | **0.001** | **4.477** | **0.000** | **[0.003, 0.008]** |
| DorsAttn | 0.002 | 0.001 | 1.567 | 0.117 | [-0.001, 0.005] |
| **SalVentAttn** | **0.004** | **0.001** | **2.655** | **0.008** | **[0.001, 0.006]** |
| **Limbic** | **0.006** | **0.003** | **2.150** | **0.032** | **[0.001, 0.012]** |
| **SomMot** | **0.004** | **0.002** | **2.075** | **0.038** | **[0.000, 0.007]** |
| Vis | -0.001 | 0.002 | -0.846 | 0.397 | [-0.005, 0.002] |
| Subcortical | 0.004 | 0.002 | 1.820 | 0.069 | [-0.000, 0.008] |

At the per-ROI level, we computed rank-order correlation between the two variables across participants and assessed statistical significance with 1,000 iterations of permutation. Similar to decomposed semantics in the last section, no ROI showed significant correlation between predictive coding and sustained attention score (two-tailed; FDR q < .05) potentially due to insufficient statistical power.

## Discussion

In this study, we investigated how neural semantic encoding during narrative listening varies with individual differences in sustained attention. By quantifying neural semantic encoding with a series of LLM embedding-based encoding models and estimating individual sustained attention scores with a connectome-based predictive model (saCPM; Rosenberg et al., 2016), we found that individuals with a neural signature of better sustained attention exhibit higher levels of latent gist compression and stronger predictive coding.

Individual differences in narrative comprehension are usually treated as noise around a group-average neural encoding score (e.g., Caucheteux et al., 2023; Tikochinski et al., 2025). The results reported here suggest that at least part of this variability is systematic and interpretable: participants predicted to have stronger sustained attention did not simply track semantic information more accurately overall, but showed a shift in what kind of semantic information their brains prioritized. These findings indicate that, rather than acting as a uniform gain on a fixed representational format, sustained attention reshapes the level of abstraction at which the brain represents an unfolding story.

Beyond its theoretic implications, the fact that encoding performance varies systematically with sustained attention scores suggests that brain-LLM alignment metrics could serve as a marker for cognitive phenotyping rather than a simple group-level index for shared representation. This extends recent work linking trait-level cognitive phenotypes to encoding fidelity (Gu et al., 2025; Vanneau et al., 2025) by showing that sustained attention relates to semantic tracking in not only its one-dimensional amplitude but also its abstraction and temporal hierarchies. Similar approaches may eventually help predict individual differences in comprehensive cognitive and behavioral metrics of interest (e.g., biases, aging, etc.). We frame this cautiously: the current study, with a single-dataset correlational approach, serves as a starting point for phenotyping applications; a validated instrument would require reliability across datasets.

An alternative interpretation of the double dissociation between low- and high-level semantic encoding is that high-level embeddings are easier to encode/decode in general (e.g., due to lower dimensionality or smoother temporal dynamics), thus inflating sensitivity to attention. We consider this less likely, as we measured encoding performance at the group level and attention sensitivity across individuals, thus a higher group mean/median encoding performance does not guarantee larger variance across individual encoding performance. On the contrary, if encoding/decoding is easier, we should expect sustained attention to have less impact on encoding performance, as every individual would achieve a high encoding score regardless.

Several limitations exist with the current study. First, sustained attention was estimated using a brain-based measure; while it is a validated measure of sustained attention trained on an independent population and task (Rosenberg et al., 2016), it is a proxy for attention rather than a direct behavioral index. Further, saCPM scores can reflect both trait-like and state-like aspects of sustained attention (Kardan et al., 2022; Rosenberg et al., 2020), which we were unable to distinguish here as the current dataset only includes one MRI session and scan type (narrative listening) per participant. Future work is needed to disentangle contributions of attentional traits and states to neural encoding performance. Second, all encoding analyses relied on GPT-2 embeddings; the layer-wise correspondence to “local" versus "gist-level" information could be specific to its architecture and training objectives resulting abstraction properties, and it is unclear how findings would generalize to other LLM families with different architectures or training regimes. Finally, the current study is correlational and uses a single narrative and dataset, precluding causal claims about sustained attention’s role and limiting generalizability across languages, narratives, and populations.

In conclusion, this study suggests that individual differences in neural encoding of narrative semantics include qualitative changes beyond the matter of a quantitative “volume knob”, enhancing gist representation but not at the expense of lower level ones; and that sustained attention may be one lever the brain uses to select a representational regime suited for predicting upcoming information. A natural next step would be to manipulate sustained attention directly (e.g., via task load or pharmacological modulation), and test whether this causally shifts the balance between local and gist-level encoding and downstream predictive coding. Extending the analyses across multiple LLM architectures, narratives, populations, and languages would help test the robustness of the compression account. Finally, applying this framework to clinical or developmental populations with known sustained attentional differences could test whether representational-format shifts, rather than raw encoding strength, better characterize variability in narrative comprehension.

Taken together, these results reframe individual variability in brain-LLM alignment from a nuisance to be averaged away into a window onto how sustained attention shapes the format, not just the strength, of semantic representation during narrative comprehension.

## Data and Code Availability

The *Le Petit Prince* fMRI dataset is publicly available at https://openneuro.org/datasets/ds003643/versions/2.0.7.

Code will be available on GitHub upon manuscript acceptance.

## Acknowledgement

We thank Ziwei Zhang for helpful discussions. We also acknowledge the use of the University of Chicago Data Science Institute computing cluster for computational resources that supported this work.

## AI Use Acknowledgement

During the preparation of this work, the authors used AI-based tools (large language models) to assist with literature search and brainstorming, as a coding assistant for data analysis scripts, and to iteratively refine the language and clarity of the manuscript text. All AI-assisted outputs—including literature suggestions, code, and written text—were iteratively reviewed, verified, and edited by the authors, who take full responsibility for the content, accuracy, and integrity of the final published work.

